# Multipoint and large volume fiber photometry with a single tapered optical fiber implant

**DOI:** 10.1101/455766

**Authors:** Filippo Pisano, Marco Pisanello, Emanuela Maglie, Antonio Balena, Leonardo Sileo, Barbara Spagnolo, Minsuk Hyun, Massimo De Vittorio, Bernardo L. Sabatini, Ferruccio Pisanello

**Author notes:** These authors contributed equally to this work (Massimo De Vittorio, Bernardo L. Sabatini, Ferruccio Pisanello).

## Abstract

Techniques to monitor functional fluorescence signal from the brain are increasingly popular in the neuroscience community. However, most implementations are based on flat cleaved optical fibers (FFs) that can only interface with shallow tissue volumes adjacent to the fiber opening. To circumvent this limitation, we exploit modal properties of tapered optical fibers (TFs) to structure light collection over the wide optically active area of the fiber taper, providing an approach to efficiently and selectively collect light from the region(s) of interest. While being less invasive than FFs, TF probes can uniformly collect light over up to 2 mm of tissue and allow for multisite photometry along the taper. Furthermore, by micro-structuring the non-planar surface of the fiber taper, collection volumes from TFs can also be engineered arbitrarily in both shape and size. Owing to the abilities offered by these probes, we envision that TFs can set a novel, powerful paradigm in optically targeting not only the deep brain, but, more in general, any biological system or organ where light collection from the deep tissues is beneficial but challenging because of tissue scattering and absorption.

## Introduction

*In vivo* fluorescence detection techniques are gaining traction in neuroscientific research as they enable recording and studying functional signals from genetically-defined neural populations from deep brain regions in freely moving animals^1^. An example is fiber photometry, an approach that allows monitoring neural activity from specific cell types in freely moving animals by recording fluorescence variations over time^2–9^. This is driving forward the development of technologies based on photonics and optoelectronic platforms^10^ as well as new methods to access multiple subpopulations using spectrally multiplexed recording of neural activity^11–13^. Typically, fiber photometry protocols rely on flat cleaved optical fibers (FFs) to stimulate and interrogate activity indicators^2–9,11–19^.

However, the accessible recording depth with flat cleaved optical fibers (FFs) is restricted to the vicinity of the fiber tip. This is due to tissue scattering and absorption effects, which, combined with the probe’s geometry, influence the fluorescence excitation and collection efficiency^20^^,^^21^. As shown by the profile in Fig. 1a (discussed in detail in ref.^20^), even when scattering and absorption effects are neglected, the amount of signal that a FF collects decreases steeply with tissue depth. In addition, reconfiguring light collection geometry to reach deeper, multiple regions requires inserting and often repositioning the fiber. Owing to the invasive geometrical profile of FFs, this unavoidably results in significant tissue damage and, in the brain, in conspicuous astrocytes and glial activation around the device even long after the implantation^22^^,^^23^. Nonetheless, FFs are widely employed to assess neural activity from deep brain regions^3^^,11–19^.

**Figure 1:**
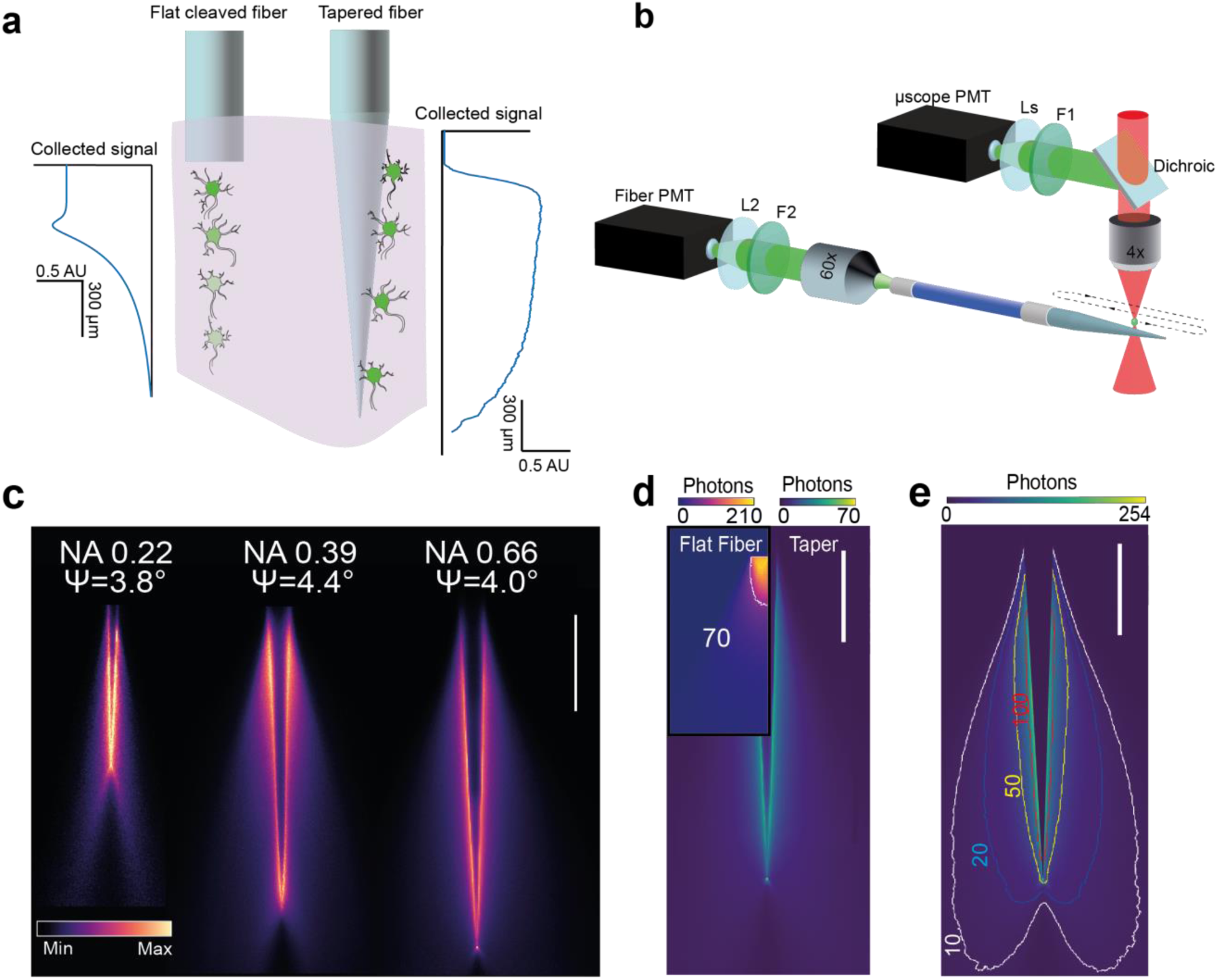
Light collection from tapered optical fibers (TFs) **a)** Schematic of light collection from a flat fiber (FF) and a TF inserted in brain tissue. The signal collected from FFs decreases steeply with tissue depth, as shown in the semi-empirical collection profile on the left inset (adapted from ref. 22). Conversely, the large optically active area of the TF collects light from a longer dorso ventral extent, as shown in the simulated profile in the right inset. In both insets, tissue scattering and absoprtion are neglected (Supplementary Fig. 1). **b**) Schematic of the optical setup (Online Methods for details). When a two-photon fluorescence spot is raster scanned around a tapered fiber, submerged in a PBS:fluorescein droplet, the generated fluorescence is simultaneously detected in epifluorescence by a non-descanned PMT (*µscope* PMT) and at the distal end of a patch fiber connected to the TF (*fiber*PMT). Abbreviations:Ls, lens system; F1, F2 bandpass fluorescence filters, L2 lens; (see Online Methods for details). **c**) Typical ξ_T_(x,y) collection fields for NA=0.22, NA=0.39 and NA=0.66 TFs in a PBS:fluorescein solution. TFs collect photons from their whole optically active region, covering depths that depend on the fiber NA, core size and taper angle. Each filed is normalized to its maximum. **d)** Comparison of number of photons collected from a two-photon fluorescence spot raster scanned in a PBS:fluorescein solution (pixel dwell time 3.2 µs) for a FF (inset) and TF with NA 0.66 and taper angle ψ~4°; the isoline in the FF inset shows the maximum number of photons collected by the TF; scale bar represents 500 µm. **e**) Isometric lines of photons collection for a NA=0.66 TF in a PBS: Fluorescein solution (30 µM); the color bar shows the number of photons per pixel (dwell time 3.2 µs); isolines are drawn at 10, 20, 50, and 100 photons; scale bar represents 500 µm.

Here we present a new approach to overcome these limitations - we exploit the modal properties of light propagation in tapered optical fibers (TFs) to structure light collection patterns on their large optically active area and to access deeper cells. In addition to be less invasive than FFs^22^, TFs probes have unique light collection features: (i) a more uniform interface over up to 2 mm of tissue along the fiber axis, (ii) the ability to perform multisite collection along the taper by time-division multiplexing, and (iii) the ability to design arbitrary collection volumes by micro-structuring the non-planar surface of the fiber taper.

Below, we first quantify the three-dimensional (3D) light collection fields of TFs, finding that TFs collect a uniform fluorescence signal from large functional regions such as the cortex or the striatum in the mouse. When combined with large area light delivery^22^^,^^24^, this property translates in a higher signal collected from TF than FF for similar illumination power density. We also show that TFs enable multipoint probing of fluorescence signal exploiting site selective light delivery. In this regard, TFs probes allow for photometry experiments that dynamically record multiple, spatially-confined brain regions at arbitrary depth along the taper. Finally, we combine modal effects governing light propagation along the taper with micro- and nano-structuring the surface of metal-coated TFs to engineer the collection volume^25,26^. This permits us to further restrict the collection volume to an angular portion of the taper surface, such that optical windows located at specific depths along the TFs can optically interface with few cell bodies in deep cortical layers. Coupled with site selective light delivery from optical windows^25^ this enables a bi-directional interface with cellular volumes at depth with high spatial selectivity.

## Results

### Light collection properties of TFs

To gather quantitative information, we first characterized the light collection properties of tapered fibers (TFs) in quasi-transparent fluorescent solution. This was done by implementing a two-photon (2P) scanning system to generate confined fluorescence spots, acting like isotropic point-like sources, in a PBS:fluorescein (30 µM) droplet in which we submerged the TFs (Fig. 1b). The fluorescence generated while raster scanning the spot around the taper was collected by two photomultiplier tubes (PMT), both synchronized with the galvo-galvo scan head: (i) the *µscope* PMT, placed in a standard non-descanned, epi-fluorescence path and (ii) the *fiber* PMT, placed at the distal end of a fiber patch connected to the TF (Fig. 1b)^20,21^. Once flattened with the reference image obtained by the *µscope* PMT, the signal from the *fiber* PMT gives access to the *fluorescence light collection field* of the TF, defined as ξ_T_(x,y). Typical collection fields ξ_T_(x,y) are shown in Fig. 1c for fibers with different numerical apertures (NA) and core diameters (NA/core diameter 0.22/50µm, 0.39/200µm and 0.66/200µm) but similar taper angle ψ~4°. We found that the light-sensitive region along the taper, defined as the collection length L, grows with increasing fiber NA and decreasing taper angle (Supplementary Fig. 2). This implies that the collection length of a TF can be tailored from a few hundreds of micrometers up to ~2 mm by modifying the fiber NA and taper angle ψ. This finding sets an important difference in collection properties between TFs and FFs, as for FFs the collection depth does not significantly depend on the NA^20^. A comparison between collection fields for TFs and FFs with NA=0.66 is shown in Fig. 1d (the same comparison for NA=0.39 is shown in Supplementary Fig. 2). The optically active surface of TFs extends along the waveguide axis, resulting in a relatively uniform collection distribution along the whole taper surface. This happens because the TF surface optically interfaces with the surrounding environment via modal subsets of increasing transverse propagation constant for wider waveguide diameters27. Therefore, the full set of propagating modes supported by the straight portion of the fiber is progressively populated along the taper section. For a FF, instead, all the propagating modes are coupled at the fiber facet. As reflected in the collection fields ξ_F_(x,y), FFs collect a higher signal intensity in the vicinity of the end facet. Conversely, TFs collection profile is maximized to a lower value in the vicinity of the taper surface and follows a two-lobe shape that widens at the tip, as shown by iso-intensity collection lines (Fig. 1e for a NA=0.66 TF, Supplementary Fig. 2 NA=0.39). In three dimensions, the distribution of the collected signal is fully symmetrical around the taper axis, as testified by volumetric collection maps obtained extending the x-y raster scanning to a *volumetric collection field* ξ(x,y,z) (Supplementary Fig. 2).

**Figure 2:**
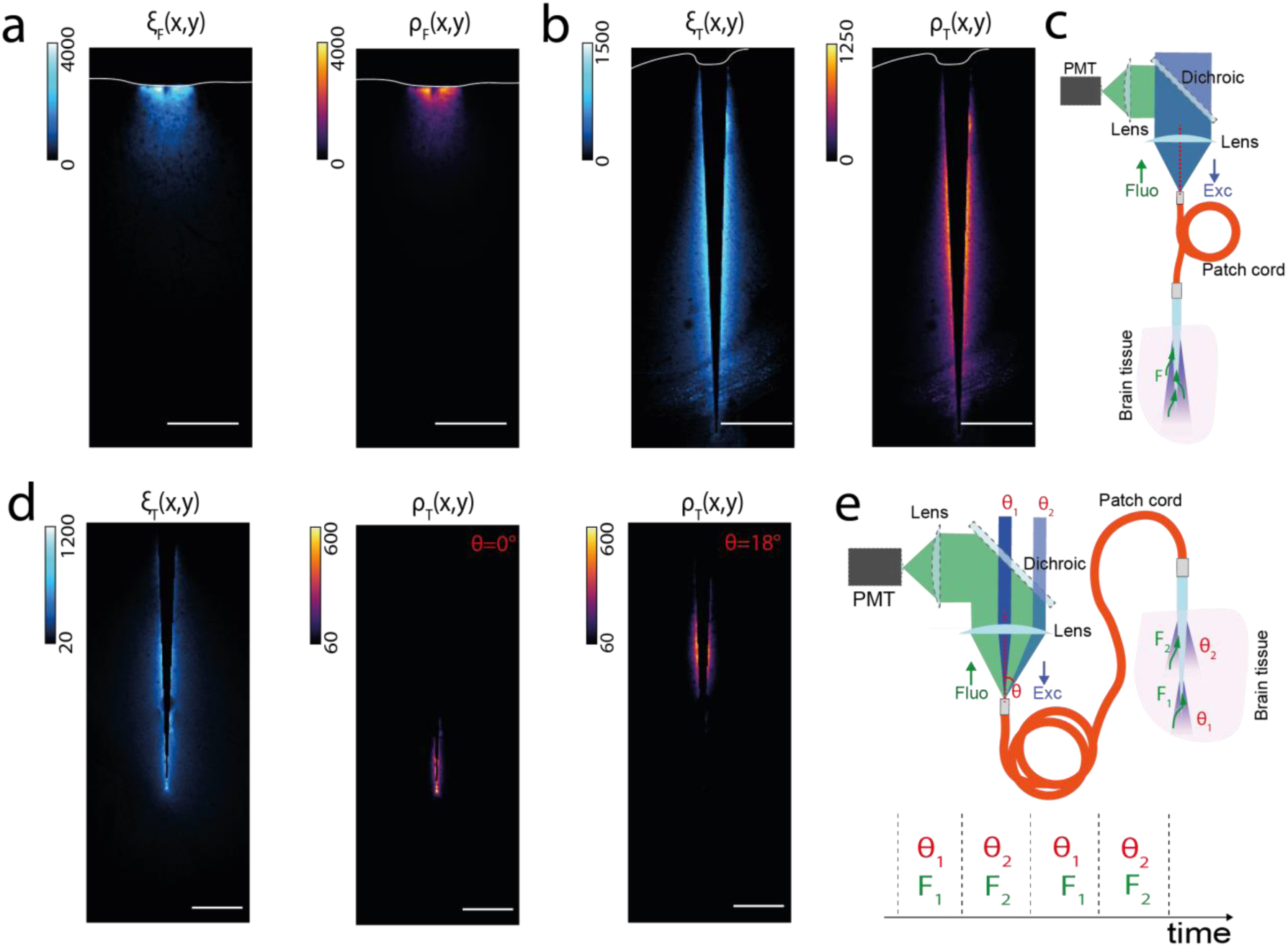
Reconfigurable light collection with TFs. **a**) Light collection field ξ(x,y) (*left*) and photometry efficiencty field ρ(x,y) (*right*) for a NA=0.66 FFs in contact with the cortex of a brain slice uniformly stained with fluorescein; the white circle indicates the center of gravity of the ξ(x,y) and ρ(x,y) fields; **b**) as in A for a NA=0.66 TF inserted in a brain slice uniformly stained with fluorescein. With wide volume illumination, the TF collects a uniform amount of signal across the whole cortex. **c**) Schematic of a system using full-NA illumination with blue laser light to stimulate and collect fluorescence from large brain region. Fluorescence is generated and detected around the whole optically active region of the taper. The collected light is back propagated in the fiber patch and discriminated from blue light by a dichroic mirror that redirects it towards a PMT. **d**) Site selective illumination, instead, allows confining the volume sampled to multiple sub-regions along the TF surface. (*Left)* Light collection field ξT(x,y), (*center*) photometry efficiency field ρT(x,y) obtained with selective illumination at the fiber tip; (*right*) ρT(x,y) field obtained via selective illumination at wider taper diameters. **e**) Schematic of a proposed system for multi-site photometry using a PMT detector in a time multiplexed configuration. A blue laser beam is launched in a fiber patch cord at increasing input angles (θ_1_, θ_2_). When injecting at low angles, laser light is outcoupled at the taper tip and generates fluorescence signal F_1_; conversely, when injected at θ_2_ laser light is outcoupled at larger taper diameters and generates fluorescence signal F_2_. The fluorescence light is detected by a PMT whose output signal is synchronized with the light injection stimulus. The fluorescence signal is attributed to the appropriate region according to its time-stamp. All scale bars represent 250 µm.

### Uniform collection over large and deep areas

Light collection properties of TFs, shown above in a semi-transparent medium, can be exploited to obtain uniform collection over large and deep brain areas in presence of scattering and absorption. To prove this, we measured *fluorescence collection fields* ξ(x,y) and *fluorescence excitation fields* β(x,y) for FFs and TFs in brain slices uniformly stained with fluorescein. These data, when combined, describe the dependence of fluorescence signal on the excitation light intensity^20^^,^^21^ by retrieving *photometry efficiency fields* ρ(x,y). These fields give exhaustive geometrical information on the sampled volume of tissue. We compared collection and photometry fields for a FF and a TF of matching NA and core size (NA=0.66, core size 200 µm). As shown by ξ_F_(x,y) and ρ_F_(x,y) fields for a FF inserted at the cortex surface (Fig. 2a), FFs effectively interface with the superficial layers of the cortex; however, they extract little information farther than 300 μm from the facet. Conversely, TFs interface more uniformly with brain tissue surrounding the optically active area of the taper, as shown by ξ_T_(x,y) and ρ_T_(x,y) fields for a TF inserted in the same brain slice (Fig. 2b). The schematic of an optical system that can be used to inject and collect light over the full taper surface is shown in Fig. 2c. Exploiting the symmetry of ξ(x,y) fields in three dimensions, we calculated the volume sampled by the waveguides as a function of the collected signal (Supplementary figure 3). Assuming a uniform neuron density of 10^5^ neurons/mm^3^ in the mouse cortex^28^, a rough estimation shows that the signal collected by TFs summarizes the contribution of more neurons than the signal arising from FFs (Supplementary figure 4). This in turn means that TFs can potentially extract more information, since the signal collected by FFs arises largely from only the reaching immediately adjacent to the open facet^20^^,^^21^ (Supplementary figure 3, 4).

### Reconfigurable multisite collection along the taper

The collection volume of TFs can be dynamically confined to multiple locations along the taper by using site-selective light delivery based on mode division de-multiplexing strategies^22^^,^^27^^,^^29^. To define the geometrical configuration of the probed volume with this approach, we first acquired the ξ_T_(x,y) collection field for a NA=0.66 TF inserted in a brain slice uniformly stained with fluorescein (Fig. 2c). Then, using a fast scanning system based on a galvanometric mirror, we injected laser light in the TF exciting modal subsets of increasing transverse propagation vector component (Supplementary Fig. 5, Online Methods). In this way we restricted the illumination volume to confined regions that can be gradually moved along the taper portion by acting on the light input angle (Supplementary Fig. 5, Online Methods)^22^^,^^27^^,^^29^. As the illumination light is delivered at specific locations, fluorescence signal is generated in a confined volume of tissue (Supplementary Fig. 5). This in turn means that TF can be employed to dynamically interrogate multiple sites across a functional region. As a proof of principle, we measured the photometry efficiency fields ρ_T_(x,y) combining the collection field ξ_T_(x,y) with emission fields β(x,y) arising from site-selective illumination (Fig.2d, Supplementary Fig. 5). As shown in Fig. 2d, the photometry signal is maximized in confined regions whose location can be promptly inferred from the light injection angle. Leveraging on this property, the fluorescence signal can be unambiguously attributed to the brain region that is receiving illumination using a time-division multiplexing system as proposed in Fig. 2e. Briefly, this can be obtained launching a laser beam in a fiber patch cord at increasing input angles (θ1, θ2). This will excite different modal subsets that are outcoupled at confined positions along the taper. The fluorescence generated for each illumination position (F1, F2 respectively) is collected by the taper, back- propagated in the fiber patch cord, discriminated by a dichroic mirror and finally detected by a PMT whose output signal is synchronized with the light injection stimulus (Fig. 2e). As shown by the photometry efficiency fields ρ_T_(x,y) (Fig. 2d), this approach is resilient to modal mixing along the patch cord^22^.

### Enhanced fluorimetry in genetically stained neural populations

To illustrate the advantages offered by the extended collection depth of TFs, we corroborated our results using a TF (NA=0.39, ψ~4°) and a FF (NA=0.39) to stimulate and detect fluorescence from different cortical layers. This was done in fixed brain slices of Thy1-ChR2-eYFP mice, where eYFP expression is limited to L2-3 and L5 (Supplementary figure 6, Online Methods). ξ(x,y), β(x,y) and ρ(x,y) fields were measured for three experimental configurations: (i) a 0.39 NA FF attached to superficial layers (Fig. 3a), (ii) a FF inserted as deep as L5 (Fig. 3b) and (iii) a TF inserted across the whole cortex extent (Fig. 3c). As shown by the ξ(x,y) and ρ(x,y) fields for the three configurations (Fig. 3), the TF stimulates and collects fluorescence from both L2-3 and L5, while the FF recruits signal within a confined region in the vicinity of their end facet and needs to be repositioned to address both areas, with consequent tissue damage. In addition, the blunt geometrical profile of FFs hinders the fiber insertion as displaced tissue remains in front of the open facet as the fiber travels down.

**Figure 3:**
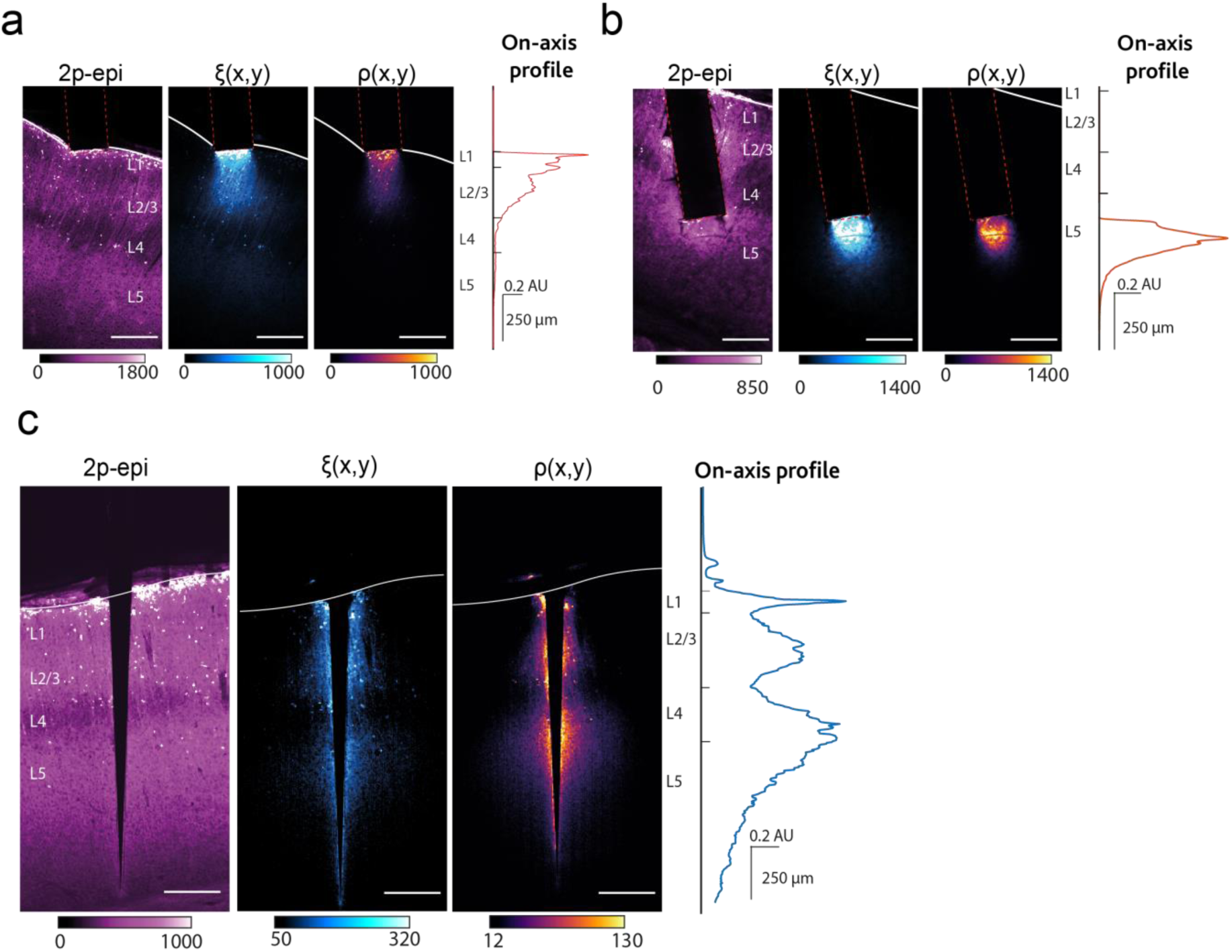
Enhanced photometry from genetically stained neural populations. **a**) Light collection for a NA=0.39 FF inserted in proximity of cortical layers L2/3; from *left* to *right*: two-photon epifluorescence image, ξ(x,y) field, ρ(x,y) field, *on-axis* collection profile extracted from ρ(x,y). **b**) As in **a** for a NA=0.39 FF inserted in proximity of cortical layers L5. In this configuration, FF insertion displaces a significant portion of brain tissue, visible from the 2p-epifluorescence image. **c**) As in A-B for a NA=0.39 TF inserted across the entire cortex. As demonstrated by the ρ(x,y) field, the TF collects and excites fluorescence from the full cortex extent. All scale bars represent 250 µm.

We compared the absolute level of signal arising from the three experimental configurations, with the fiber probes delivering blue laser light (473 nm) while recording the collected fluorescence in the 500-550 nm band with a PMT (Fig. 4a). When using the same average power density at the optical surfaces of FFs and TFs (~0.1 mW/mm^2^), the TF collects a larger fluorescence signal with respect to the FF at both depths (Fig. 4b). This can be explained considering the combined effect of mode division de-multiplexing in light collection and light delivery. At first, as argued above, the integrated signal collected by TFs receives a significant contribution from a higher number of neurons (Supplementary Fig. 4). In addition, mode-division de-multiplexing distributes the higher illumination power over a wider surface, while maintaining a moderate power density^22^^,^^27^. As more photons are released into the tissue and more neurons contribute to the collection signal, a higher level of fluorescence is generated and detected. This observation is consistent with previous results finding that TFs can elicit optogenetic activation with smaller output powers than FFs in the same experimental model^22^. Importantly, since photobleaching is dependent on the exposure of each fluorophore to photons, the more efficient distribution of light by TFs allows more fluorescence to be generated without increasing photobleaching.

**Figure 4:**
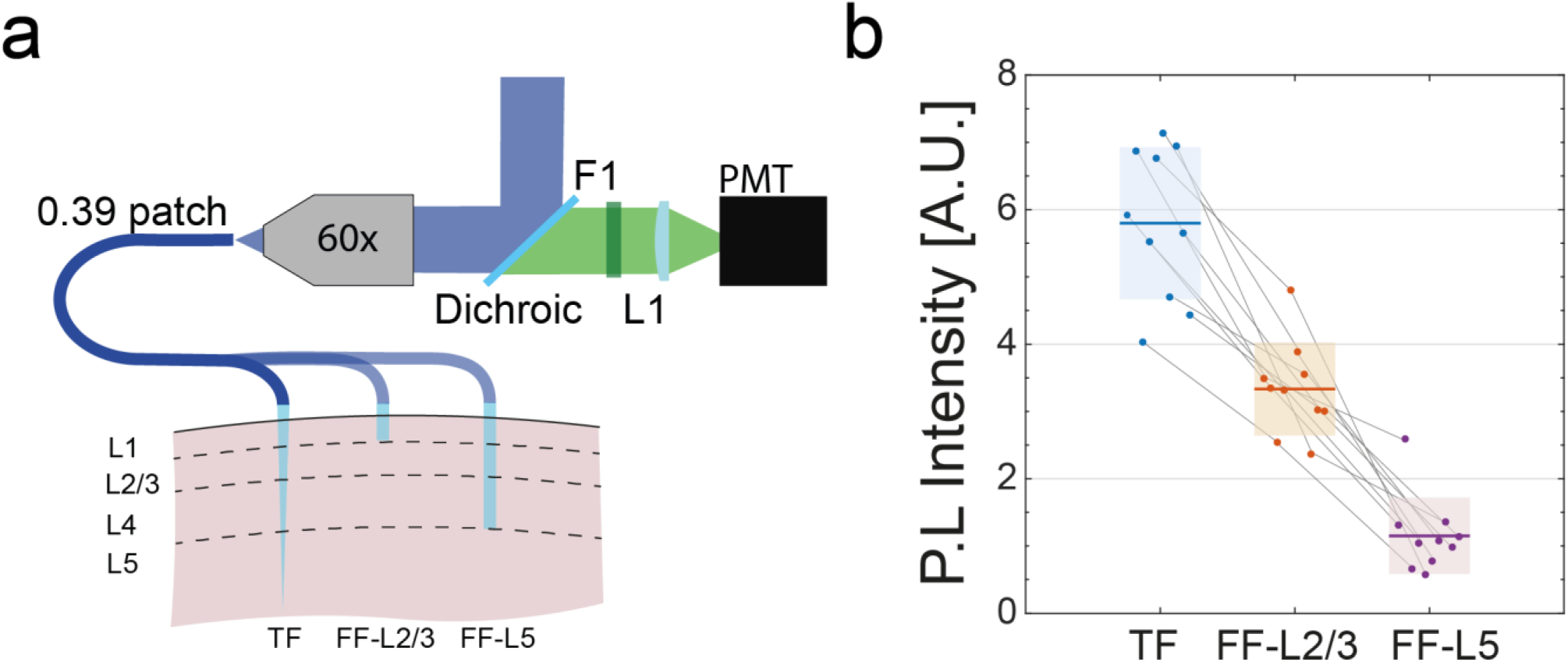
Extracting enhanced information from large and deep brain regions. **a**) Diagram of the photometry system for the three experimental configurations. Blue laser light (473 nm) is injected into a fiber patch that couples with the waveguides: a TF inserted across the whole cortex, a FF inserted in layer 2/3, a FF inserted as deep as layer 5. **b**) Intensity of the fluorimetry signal (N=10) in slices from Thy1-ChR2-YFP mice using 0.39 TFs (ψ=4°) inserted across the cortex (blue), a 0.39 FF inserted in L2/3 (orange) and 0.39 FF in L5 (purple) to stimulate and detect fluorescence (see Methods). Laser power was adjusted to obtain a similar power density from the optically active area of FF and TFs (0.1 mW/mm^2^ at 473 nm). Shaded areas indicate data standard error on the mean for each experimental configuration. Grey lines connect data obtained in the same experimental run from the same brain slice. Tapered fiber consistently collect a higher signal than FFs in both configuration. Statistical significance was assessed with a Student *t* test with significance α=0.001 (see Online Methods).

### Reconfigurable multisite collection from genetically stained neural populations in the mouse striatum

To gain a deeper understanding of neural processes that underlie motor behavior and the regulation of reward driven actions, recent works adopted fiber photometry techniques to optically interface with striatal neurons^11–13,^^30^. In this context, TFs might add experimental capabilities by sampling multiple regions with a single, remotely controlled implant. To support this argument, we characterized site-selective fluorimetry sampling with TF in the striatum. A NA=0.66 TF was inserted in the striatum region of fixed brain slices from Thy1-eYFP mouse and, after acquiring the ξ(x,y) field (Fig. 5b), site-selective illumination was used to produce photometry efficiency ρ(x,y) fields for increasing input angles (Supplementary Fig. 5, Online Methods). As shown in (Fig. 5c), the sample volume that responds to site-selective illumination gradually moves away from the TF tip as the light input angle is increased.

**Figure 5:**
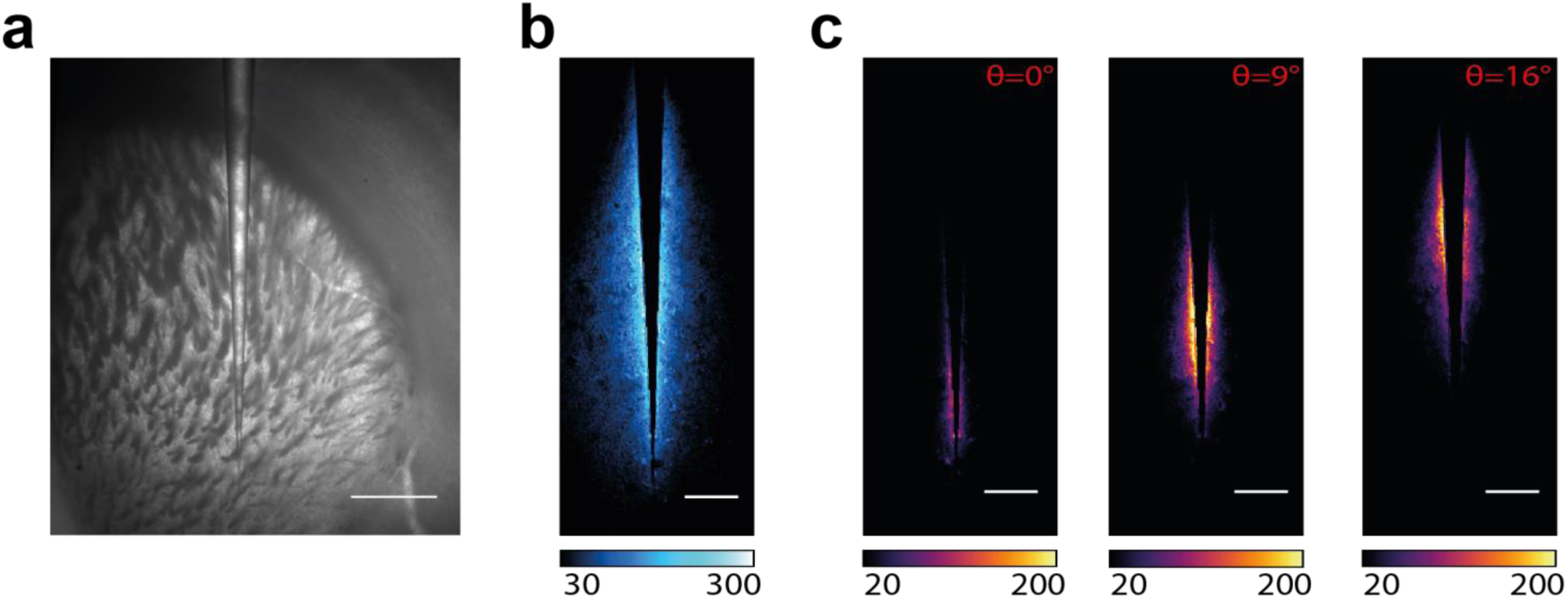
Multisite photometry sampling of genetically targeted neurons in the mouse striatum. **a**) Brightfield image of a TF inserted in the striatum region in a fixed brain slice from Thy1-eYFP mouse. **b**) Light collection field ξ_T_(x,y) for the TF shown in panel A. **c**) The combinaton of the ξ_T_(x,y) field (a) with site-selective light delivery, generates photometry efficiency fields ρ_T_(x,y) that show reconfigurable, multisite light collection from sub-regions in the striatum. Scale bars represent: 500 µm in panel a and 250 µm in panel b and c.

### Engineering collection volumes with micro-structured TFs

The availability of a large surface that can interface with the environment confers to TFs the possibility of engineering the collection volume according to the region of interest. This can be obtained using micro and nano-fabrication technologies to structure the fiber taper, as previously demonstrated for light delivery^25^^,^^26^^,^^31^. Examples of this approach are given in Figures 5 and 6, where we demonstrate: (i) the restriction of light collection to a specific angular extent around the waveguide (Fig. 6), (ii) the observation of cell bodies in deep layers of the cortex while restricting light collection to a specific spot along the taper (Fig. 7).

**Figure 6:**
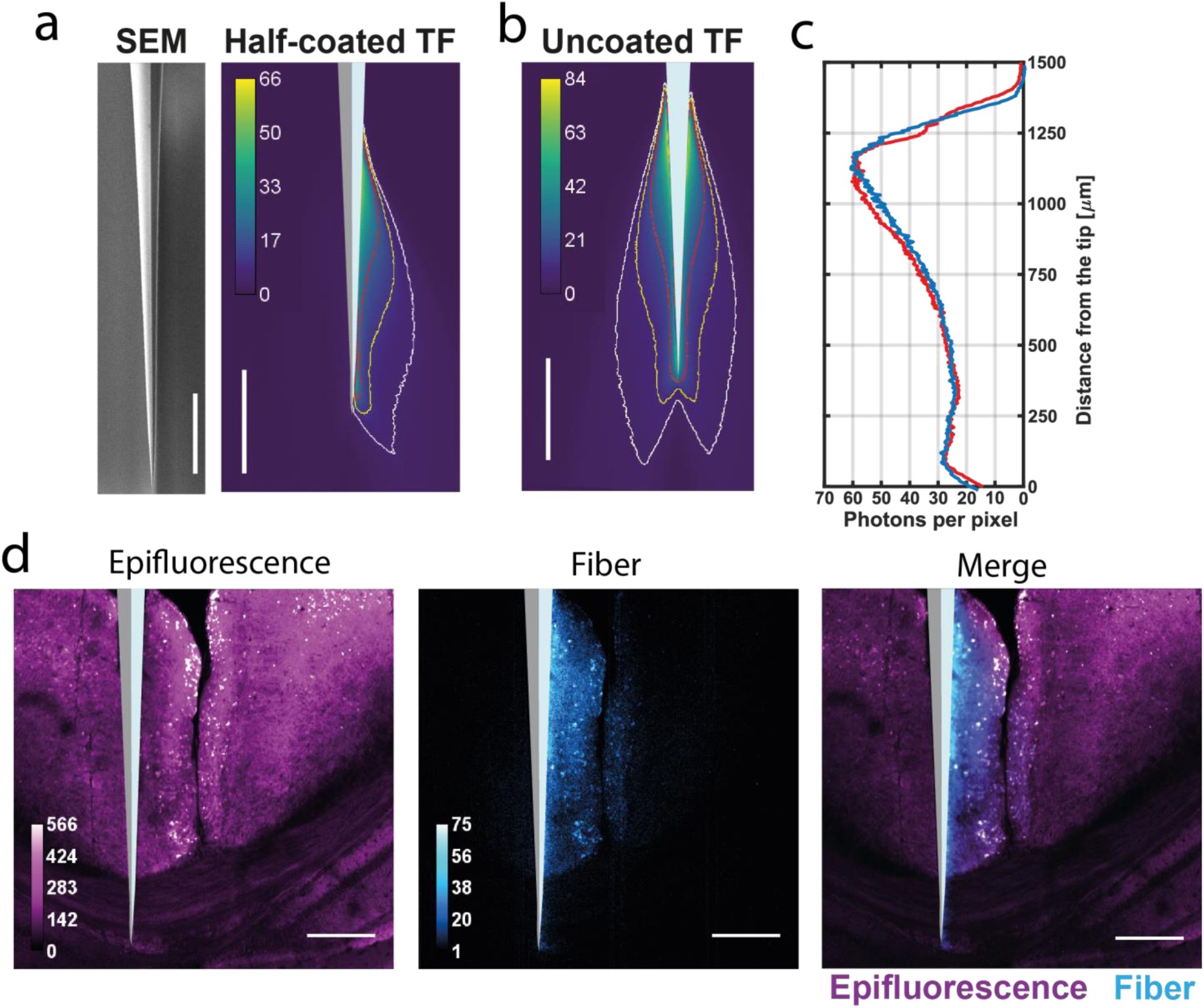
Side selective light collection from half-coated TFs. **a**) *left*, representative SEM micrograph of a TF half-coated with Al; *right*, ξ(x,y) field for a half coated NA=0.39 TF in PBS:fluorescein solution (30 µM); an overlayed diagram of the half-coated probe shows the orientation of the uncoated surface; **b**) ξ(x,y) field for an uncoated NA=0.39 TF in PBS:fluorescein solution (30 µM); **c**) Intensity profiles extracted from ξ(x,y) fields for a half-coated (red) and an uncoated TF (blue). **d**) Side-selective light collection from a half-coated TF inserted in a fixed brain slice from Thy1-ChR2-YFP mouse strain; from left to right: two-photon epifluorescence image, ξ(x,y) field, merge of epifluorescence (magenta) and ξ(x,y) field (cyan). All scale bars are 250 µm.

**Figure 7:**
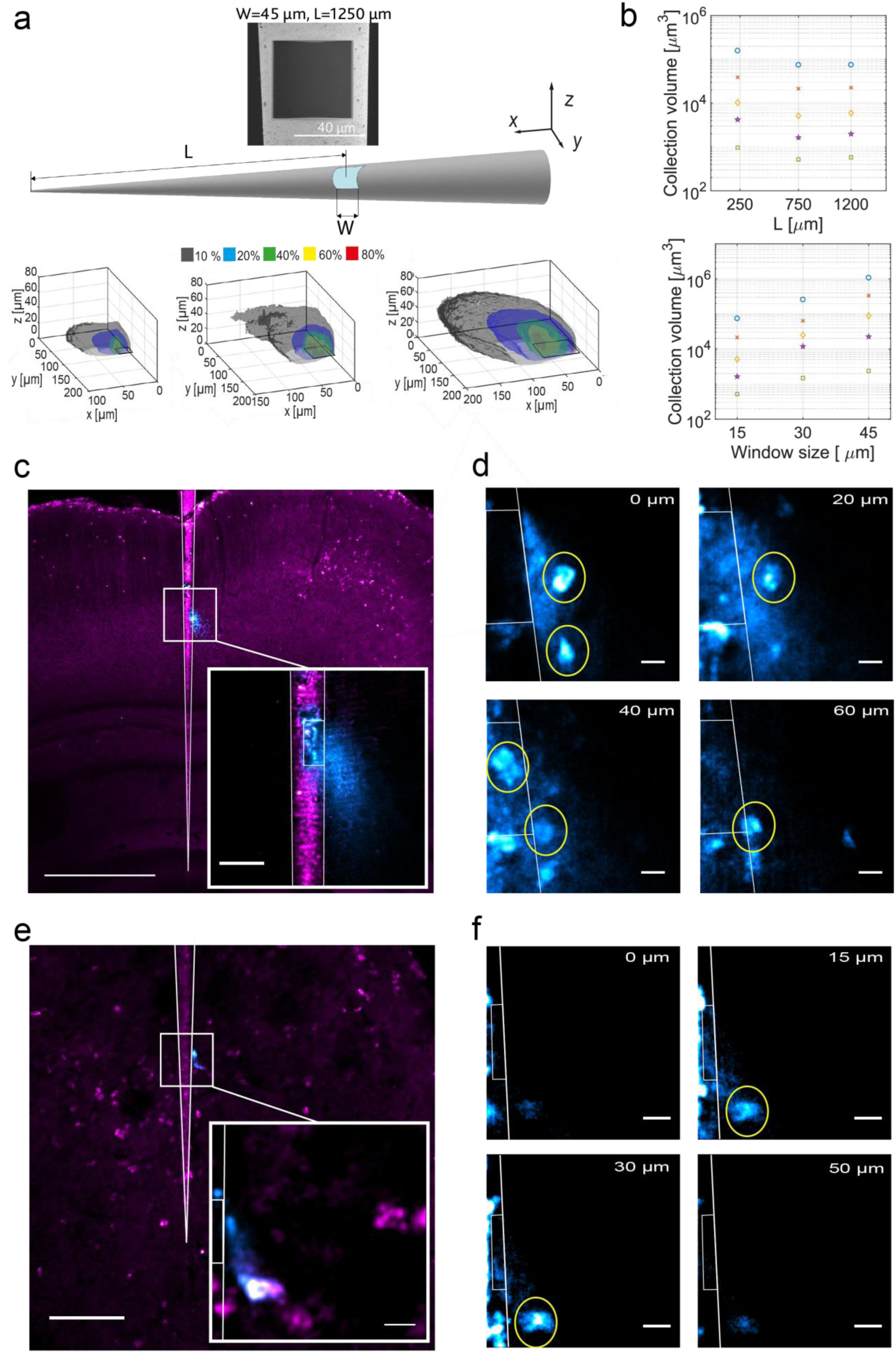
Engineering light collection volume with micro-structured TFs. **a**) *top*, diagram of a NA=0.39 micro-structured TF with a squared optical window of size W opened at distace L from the fiber tip; *inset*, SEM micrograph of a W~45 µm square optical window opened at L~1250 µm from the fiber tip; *bottom*, 3D maps of ligth collection obtained from windows of different sizes in a PBS:fluorescein solution (30 µM); isosurfaces of collection are drawn as fraction of the maximum number of collected photons (10%, 20%,40%,60%,80%). **b**) *top*, collection volumes enclosed by collection isosurfaces versus window distance from the tip (L=250, 750, 1200 µm); *bottom,* collection volumes enclosed by collection isosurfaces versus window size (W=15, 30 60 µm). **c**) Light collection from a micro-structured TF with a W~45 µm square window at L~750 µm from the tip inserted in a fixed brain slices from Thy1-ChR2-YFP mouse strain; the image shows the overlay of the ξ(x,y) filed (cyan) with a simultaneously acquired two-photon epifluorescence image (magenta); scale bar represents 500 µm; *inset*, a close up on the window shows a confined collection volume, scale bar is 50 µm; **d**) frames from a z-stack at increasing height above the optical window (0 µm, 20 µm, 40 µm, 60 µm) demonstrate the ability of optical sectioning with single cell sensitivity when acquiring fluorescence through the optical window; the TF profile and the window position are outlined in white; scale bar is 10 µm. **e**) as in panel c for a TF with a W~20 µm square window at 230 µm from the TF tip; scale bars represent 100 µm for the large view and 10 µm for the *inset*. **f**) as in D for a W~20 µm optical window: frames from a z stack at increasing height (0 µm, 15 µm, 30 µm, 50 µm) shows optical sectioning with single cell sensitivity; the TF profile and the window position are outlined in white; scale bar is 10 µm.

To confine light-collection to a half-portion of the waveguide surface, we thermally evaporated a highly-reflective Al layer on the opposite side (Fig. 6a, see Online Methods for details on the evaporation process). The resulting collection field ξ(x,y) is compared with the same measurement for a twin (uncoated) TF (Fig. 6b). The collection profile along the optical surface of the half-coated TF is very similar to the profile recorded by the uncoated probe, both in terms of shape and of collected number of photons (Fig. 6c). Therefore, the metal coating does not substantially change the detected signal, letting us suggest that almost all photons entering into the uncoated taper undergo dielectric total internal reflection, a condition that is instead forced by the metal layer in the half-coated fiber. Following these considerations, we suppose that the light out-coupling mechanism of TFs^22^^,^^27^^,^^29^^,^^32^ might be inverted to describe light collection from TFs. Subsets of guided modes with low transverse propagation constant (k_t_) entering in the tip region will encounter progressively larger waveguide diameters; as the transverse propagation constant is inversely proportional to the waveguide diameter^27^^,29,32^, these modal subsets are more likely to be propagated than out-coupled as they travel upwards in the tapered region. The side-collection properties of the device were tested in brain tissue with the half-coated TF inserted in a fixed brain slice expressing YFP following the promoter Thy1 (Fig. 6c, see Methods). A 2P epifluorescence image and the ξ(x,y) field was acquired simultaneously. As expected, despite generating fluorescence all around the fiber, only fluorescence that arises close to the uncoated portion of the fiber was acquired by the *fiber* PMT (Fig.1, Fig. 6d).

### Light collection from optical windows at arbitrary depth

Building on these results, we explored the possibility of further restricting the collection volume by fabricating optical windows on a TF entirely coated with Aluminum. This was done using focused ion beam (FIB) milling to selectively remove the metal in specific regions of the taper^25^^,^^26^. To optimize the devices, light collection from optical windows in PBS:fluorescein solution (30 µM) was characterized following the methods described above. We fabricated probes (NA=0.39 TFs) with squared optical windows of different side length (W=60 µm, 30 µm and 15 µm) positioned at different distances from the fiber tip (L=230 µm, 750 µm, and 1250 µm) (Fig. 7a). For each of these devices, we acquired volumetric collection diagrams (Fig. 7a), observing that larger windows’ area lead to higher collection volume (Fig. 7b). On the other hand, while a window positioned towards the tip tends to collect from a slightly larger volume, windows at larger taper sections have similar collection properties (Fig. 7b).

Micro-structured TFs were tested in fixed brain slices from a Thy1-eYFP mouse strain. We acquired ξ(x,y) fields that we correlated to simultaneous epi-fluorescence imaging. Results reported in Fig. 7c, for a TF with a window of width 45µm and length 750 µm, show that light collection co-localizes with the optical window location (Online Methods). This property ties in well with selective light delivery from optical windows^25^ as it allows interfacing with cellular volumes at depth with high spatial selectivity. Fig.7d displays a set of images from a stack acquired from a region adjacent to an optical window with side of 45µm located at 750µm from the tip (Supplementary Fig. 6, Online Methods). The signal from identified cell bodies is clearly detectable and allows a 3D reconstruction of single cell somas from the TFs (Fig. 7d), with an exact match between photometry diagrams and epi-fluorescence imaging (Supplementary Fig. 6). Similar results were obtained using a TF with windows of 25 µm width, located at 230 µm from the tip, as shown in Fig. 7e. As for the larger window, we observed that light collection co-localizes with the window position (Fig. 7f) and we reconstructed a 3D stack for a single cell in the vicinity of the window.

## Discussion

Acquiring fluorescence signals from the brain is a powerful technique in neuroscience^33^^,^^34^ and the field would greatly benefit of an implantable waveguide system that can be configured to efficiently and selectively collect light from the region(s) of interest. In this paper we demonstrate that TFs can accomplish this goal by exploiting the modal properties of the narrowing waveguide. As the taper narrows, guided modes are gradually coupled with the environment enabling the probe to collect light along an extended region with a more homogenous collection profile compared to standard FFs (Fig. 2 a, b, Fig. 3, 4). Given a constant output power density at the active optical surface, FFs are better suited to collect signal from shallow brain regions adjacent to the fiber facet^20^. Conversely, TFs interface with a higher number of neurons distributed over a large brain region, thereby recruiting more information while mitigating tissue damage.

The modal distribution along the taper can also be exploited to perform site-selective fluorescence collection. By selecting the subset of guided modes injected into the fiber, fluorescence can be generated only around at a specific portion of the taper (Fig. 2c and 5). This can be used to inspection multiple points along the taper as light emissions is switched to different sections in a time-division fashion (Fig. 2e). We have previously demonstrated the compatibility of site-selective illumination in free-moving animals^22^.

To further tailor the shape and size of the collection volume, the highly-curved taper surface can be coated with metal and then patterned using a Focused Ion Beam. This led us to restrict the collection volume to an angular portion of the TFs (Fig. 6) and to precisely modulate the position and size of the brain region receiving from which fluorescence signal is generated and collected (Fig. 7).

TFs therefore add beneficial features to the existing set of devices for light collection from the brain, providing a set of configurations that is not achievable either with FFs or with µLED/photodetectors systems^10^. Current existing technologies can be wirelessly driven, but have a much smaller configuration versatility with respect to TFs, that can be also structured all around the taper^26,31^. This provides the ability to generate nearly arbitrary light collection patterns and therefore opens new perspectives for applications needing depth resolved information for structural, functional and diagnostic analysis of the living brain. On the strength of these results, we envision that the application of TFs probes to fluorescence collection from scattering tissue will facilitate dissecting the contribution of multiple functional areas in deep brain regions while providing for a versatile complement to existing optical methods

In evaluating new applications of TFs in experimental systems other than the brain, it will be interesting to consider that the performance shown above are based on mode-division de-multiplexing in TFs. This is an intrinsic property of tapered waveguides that is generated by the effect of the narrowing diameter on the transverse propagation constant of a given modal subset ^22^^,^^27^^,^^29^^,^^32^. The optical properties of the medium surrounding the waveguide play a role in selecting which modal subset is outcoupled or collected at a specific taper section. However, mode-division de-multiplexing properties are preserved across the ensemble of biological tissues in living animals, a class where the brain represents one of the worst cases in terms of scattering properties, considering both scattering length and anisotropy^35^.

## Materials and Methods

### Optical setup

A custom two-photon laser scanning microscope was implemented to characterize TFs light collection fields ξ (x,y,z) and light emission fields β(x,y,z). The apparatus is similar to previously described systems^20,21^. Briefly, the optical setup consisted of a scanning path followed by an upright microscope. A femto-second pulsed near-infrared laser beam (Chameleon Discovery, Coherent) was directed towards a power modulation system composed of a polarizer, a polarizing cube and a Pockel Cell (Conoptics 350-80-02). Then, a periscope raised the laser beam and a quarter wave-plate (Thorlabs AQWP05M-980) produced a circular polarization. A beam expander enlarged the laser beam before relaying it into a galvanometric scan system (Sutter). The scanned beam was then reflected towards the back aperture of an objective mounted on an upright Olympus IX-71 microscope aligned with the scanning path. A piezoelectric motor (Phisik Instrument P-725.4CD) was used to control the objective *z* position allowing for volumetric scanning. An epifluorescence detection system was set using a dichroic mirror (Semrock FF665-Di02) positioned above the objective to reflect fluorescent light towards a non-descanned PMT referred to as “µscope” PMT (Hamamatsu H10770PA-40). Light was focused on the PMT window with a lens system and filtered (BPF, Semrock FF01-520/70-25). Image acquisition was controlled via commercial software (ScanImage, Vidrio Technologies). Images were acquired in 512×512 pixels with 3.2 µs pixel dwell time. Laser power and scanning range were adjusted at convenience. Wide-field images (Fig.1, 2, 3, 5, 6) were acquired with a 4× objective (Olympus XLFluor 4x/340a); high-resolution images (Fig. 7) were obtained with a 25× objective (Olympus XLPLN25XWMP2). When necessary, image stitching was performed with FIJI software^36^.

### Light collection fields ξ (x,y)

Light collection fields were measured for TFs and FFs submerged in a semi-transparent solution (PBS:fluorescein 30 µM) or inserted in brain slices. An endoscopic detection system was implemented to detected fluorescent light collected by the fibers when the focal spot was raster scanned in their surroundings. To do this, the fibers were connected to a fiber patch via ferrule-to-ferrule butt coupling. Light emitted at the distal end of the fiber was collected with a high-NA objective (Olympus Plan N 40x). The output of the objective was routed to a second PMT referred as “fiber” PMT (Hamamatsu H7422P-40) after passing through two spherical lenses (Thorlabs LA1050-A and LA1805-A) and a band pass filter (Semrock FF03-525/50-25). Collection fields ξ(x,y) were obtained by averaging out 60 acquisitions. Volumetric collection fields ξ(x,y,z) were produced by recording the fiber output during volumetric scanning. Light collection fields in brain tissue were acquired by inserting the TF in brain slices positioned in the sample plane. Fiber insertion was controlled with a micro-manipulator (Scientifica) upon live imaging with a CCD camera. Light collection fields from optical windows in fluorescein and brain slices were obtained scanning the two-photon fluorescent spot with a high magnification 25× objective (Olympus XLPLN25XWMP2) in the vicinity of the window.

### Light emission fields β(x,y)

To measure light emission fields β(x,y,z), laser light (473 nm) was injected at the distal end of the fiber patch. For full NA light emission^22^^,^^27^, light was coupled across the whole NA of the fiber patch using a high-NA microscope objective (Olympus Plan N 40x) (Fig. 4a); for site-selective illumination, light was injected using a scanning system based on a galvanometric mirror (Supplementary Fig. 5)^22^^,^^27^. The optical system was equipped with two detection paths to image the light emission field, primarily in brain tissue.

As the optical system houses a confocal scanning path alongside the two-photon path, the detection was rearranged to direct fluorescent light excited by the fiber emission towards a descanned pinhole system (Thorlabs MPH16). This was done where bleaching effects had limited effect, such in the case of fluorescein staining. Light passing through the pinhole aperture was detected by a PMT, the “pinhole PMT”, synchronized with the galvo scanning. This approach has the advantage of providing with emission fields whose photon-to-pixel assignation intrinsically replies the one used for ξ (x,y,z) fields, thus avoiding rescaling and interpolation in image registration. However, the image formed by the objective needs to be scanned several times across the pinhole. This increases illumination time and potentially results in fluorescence bleaching.

In genetically stained brain slices, therefore, the emission fields were acquired by redirecting the dichroic mirror towards a CCD camera in bright-field mode. While this approach needs image rescaling to align emission and collection fields, is advantageous to avoid bleaching as all pixels are acquired simultaneously.

### Photometry efficiency fields ρ (x,y)

Photometry efficiency fields were calculated by scaling the collection field ξ (x,y) against the normalized illumination field β(x,y). Following the approach outlined in ref., the photometry field was calculated as

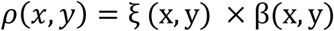

where × indicates pixel by pixel multiplication. ξ (x,y) and β(x,y) fields were registered manually using FIJI software. The inside portion of the TF was masked as the information arising from it might suffer aberrations due to the tissue-glass interface.

### Tapered fiber fabrication

Tapered fiber stubs fabricated from fiber cords with NA = 0.22 core/cladding = 50/125 μm (Thorlabs FG050UGA), NA = 0.39 core/cladding = 200/225 μm (Thorlabs FT200UMT), NA = 0.66 core/cladding = 200/230 μm (Plexon PlexBright High Performace patch cable) were obtained from OptogeniX (www.optogenix.com). Details of the fabrication procedure are provided in previous works^22^^,^^37^. Fibers were connected to metallic ferrules to minimize auto-fluorescence effects with a diameter of 1.25 mm. Half-coated fibers were fabricated by thermally evaporating 600-800 nm of Al on one side of the tapered section. This was done by exposing only one side of the fiber to the crucible. Optical windows were realized with Focused Ion Beam (FIB) milling on TF entirely coated with Al. Briefly, a uniform Al layer was deposited on the TF via thermal evaporation. To obtain a uniform coating, the fibers were mounted on a rotating motor. Optical windows of different sizes were fabricated by removing the metal coating at specific taper diameter using a dual beam FIB-SEM system (FEI Helios Nanolab 600i Dual Beam). In order to remove only the metal coating, without affecting the glass, FIB milling was supervised via simultaneous Scanning Electron Microscopy (SEM) imaging.

### Brain slices

Brain slices were obtained from wild-type mice and Thy1-ChR2-eYFP mice. Mouse brains were fixed in 4% PFA and cut with a vibratome in 300 µm-thick slices. Slices from wild type were permeabilized in 0.3% Triton X (Sigma Aldrich) for 30 min, washed in PBS (three 5 min washings), incubated in 1 mM fluorescein solution overnight and washed again (three 5 min washings). All animals were cared for and used in accordance with guidelines of the U.S. Public Health Service Policy on Humane Care and Use of Laboratory Animals and the NIH Guide for the Care and Use of Laboratory Animals.

### Fluorimetry measurements in genetically stained neural populations

Blue laser light (473 nm, Laser Quantum Ciel) was injected over the full NA of a 0.39 NA fiber patch using a microscope onjective (Olympus Plan N 40x). The patch was connected to FFs and TFs stubs via ferrule-to-ferrule butt coupling. These stubs were inserted in the brain slices using a micromanipulator (Scientifica) with the guidance of simultaneous two-photon imaging. The intensity of the fluorescence signal emitted by the brain slice was monitored with two-photon imaging to ensure that all measurements were performed in similar conditions.

Fluorimetry measurements were performed by exciting and collecting fluorescence signal from genetically stained neural population via the FFs and TFs stubs. The collected fluorescence signal was discriminated with a dichroic mirror, a fluorescence filter (Semrock FF03-525/50-25) and detected with the fiber PMT. The PMT signal was amplified and digitized with a data acquisition board (NiDAQ USB-6363) that was controlled with custom software (LabView).

In order to obtain a similar output power density from TF and FFs across all measurements, ρ_P,flat_ = ρ_P,taper_ = ρ_P_ = 0,1 mW/mm^2^,laser power was modulated according to the fiber output surface. Power density ρ_P,flat_ for FFs was calculated as

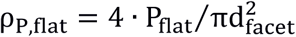

where P_flat_ is the input power and d_facet_ is the fiaber diameter (200 µm).

Power density ρ_P,taper_for TFs was instead calculated as

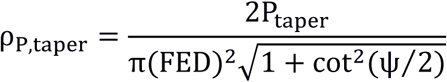

where P_taper_ is the input power, FED is the first diameter of emission (125 µm), ψ is the taper angle and clear meaning of the subscripts.

We performed 10 measurements for each of the three experimental configurations: FF in L2/3, FF in LV, TF across whole cortex, Fig. 4a. Fluorescence signal was acquired for 10 s, at 1000 samples/s, and binned in 100 ms intervals. Fig. 4b shows the mean value of the three acquisitions for each measurement; error bars are smaller than the data point size.

This produced a population of ten signal values for the three different configurations, from here, referred as T, F_placed_, F_in_ respectively for Taper, Flat placed and Flat inserted).

In order to quantitatively compare the total signal collected from the tapered fibers with the one collected by the flat fibers (at the same illumination power density) we performed a two-tailed *t* test between the two hypotheses:

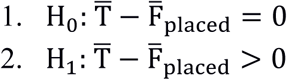

where are 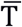, 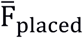 are the mean values for the respective populations. We built a statistic variable 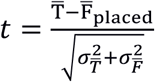 where 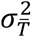, 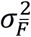 are standard errors on the mean values, obtaining t=5.88. We rejected the null hypothesis H_0_ with a significance level α = 0,001 with 18 degrees of freedom. The same procedure was applied to 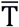 and 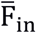 obtaining t=11.62 and rejecting H_0_ with a significance level α = 0,001 with 18 degrees of freedom.

## Acknowledgements

F.Pisano, E.M, A.B., E.M, B.S. and F.Pisanello acknowledge funding from the European Research Council under the European Union’s Horizon 2020 research and innovation program (#677683); M.P and M.D.V. acknowledge funding from the European Research Council under the European Union’s Horizon 2020 research and innovation program (#692943). L. S., M.D.V. and B.L.S. are funded by the US National Institutes of Health (U01NS094190).

